# Foot shock stress generates persistent widespread hypersensitivity and anhedonic behavior in an anxiety-prone strain of mice

**DOI:** 10.1101/700310

**Authors:** Pau Yen Wu, Xiaofang Yang, Douglas E. Wright, Julie A. Christianson

## Abstract

A significant subset of patients with urologic chronic pelvic pain syndrome (UCPPS) suffer from widespread, as well as pelvic, pain and experience mood-related disorders, including anxiety, depression, and panic disorder. Stress is a commonly-reported trigger for symptom onset and exacerbation within these patients. The link between stress and pain is thought to arise, in part, from the hypothalamic-pituitary-adrenal (HPA) axis, which regulates the response to stress and can influence the perception of pain. Previous studies have shown that stress exposure in anxiety-prone rats can induce both pelvic and widespread hypersensitivity. Here, we exposed female A/J mice, an anxiety-prone inbred murine strain, to 10 days of foot shock stress to determine stress-induced effects on sensitivity, anhedonia, and HPA axis regulation and output in. At 1- and 28-days post-foot shock, A/J mice displayed significantly increased bladder sensitivity and hind paw mechanical allodynia. They also displayed anhedonic behavior, measured as reduced nest building scores and a decrease in sucrose preference during the 10-day foot shock exposure. Serum corticosterone was significantly increased at 1-day post-foot shock and bladder mast cell degranulation rates were similarly high in both sham- and shock-exposed mice. Bladder cytokine and growth factor mRNA levels indicated a persistent shift toward a pro-inflammatory environment following foot shock exposure. Together, these data suggest that chronic stress exposure in an anxiety-prone mouse strain may provide a useful translational model for understanding mechanisms that contribute to widespreadness of pain and increased comorbidity in a subset of UCPPS patients.

## 1. Introduction

According to the American Psychological Association, nearly three-quarters of American adults in 2017-2018 stated they experienced at least one stress-related symptom over the past month [1]. This percentage jumps to 91% for Gen Z adults between 18 and 21 years of age. Stress is a well-known trigger for most chronic pain disorders, causing an onset or exacerbation of ongoing symptoms [10; 22]. Conversely, stress-related psychological disorders, such as depression, anxiety, and panic disorder, are more prevalent among chronic pain patients [8; 14; 17; 32; 37]. In patients with urologic chronic pelvic pain syndromes (UCPPS), the severity and widespreadness of pain is associated with psychological disturbances [29]. A study by Lai et al., [29] reported that patients with interstitial cystitis/painful bladder syndrome (IC/PBS) and/or chronic prostatitis/chronic pelvic pain syndrome (CP/CPPS) with numerous pain locations had worsened pelvic and non-pelvic pain severity, poorer sleep quality, greater depression, anxiety, psychological stress, and higher negative affect scores compared to IC/PBS and CP/CPPS patients with only pelvic pain.

Many patients with comorbid chronic pain and mood disorders display either hyper- or hypocortisolism, indicative of an improperly functioning hypothalamic-pituitary-adrenal (HPA) axis [15; 54]. The hypothalamus responds to stress by releasing corticotropin-releasing factor (CRF), which causes a downstream release of cortisol in humans or corticosterone (CORT) in rodents [20; 53]. Peripheral CRF and CORT release impacts both metabolic functions and neuroimmune interactions that can drive inflammation and increase pain sensitivity [15]. Higher limbic structures, including the amygdala and hippocampus, have been shown to positively and negatively regulate the HPA axis, respectively [20]. Interestingly, brain imaging studies have revealed that UCPPS patients with widespread pain and increased comorbidities have altered functional connectivity and metabolite concentrations in limbic structures associated with HPA axis output, supporting a role for altered central processing in chronic pelvic pain syndromes associated with stress [19; 27].

We have previously shown that early life stress exposure negatively impacts the HPA axis, increases urogenital and hind paw sensitivity, and evokes mast cell degranulation in male and female mice, all of which is exacerbated following adult stress exposure [16; 42; 44]. Other groups have shown that both anxiety-prone [30; 47] and normative [48] rat strains display urogenital hypersensitivity following repeated stress exposure. Here, we have adapted these previous studies to the anxiety-prone A/J mouse strain to provide evidence that exposure to repeated foot shock leads to persistent bladder and hind paw hypersensitivity, anhedonia, and altered gene expression indicative of dysregulation of limbic control of the HPA axis.

## 2. Methods

### 2.1 Animals

All experiments in this study were performed on adult (>12-week-old) female A/J mice (Stock No: 000646, Jackson lab, Bar Harbor ME USA) housed in the Research Support Facility at the University of Kansas Medical Center. Mice were housed in a climate-controlled room on a 12-hour dark-light cycle and received water and food ad libitum. Animal use protocols conformed to NIH guidelines and were approved by the University of Kansas Medical Center Institutional Animal Care and Use Committee and the Committee for Research and Ethical Issues of IASP.

### 2.2 Foot shock stress exposure

Mice were exposed to foot shock stress for 10 consecutive days according to the following paradigm. Mice were transferred to a sound-proof room and placed 4 at a time into a Tru Scan Arena system cage equipped with a shock floor (26 × 26 × 39 cm, Coulbourn Instruments Holliston, MA, USA). Mice in the shock cohort were exposed to one of five random foot shock sequences consisting of 30 0.4 mA shocks over a 15-minute period. Mice in the sham cohort were held in the Tru Scan Arena system cage for 15 minutes, but received no shocks. Mice were returned to their home cages after foot shock or sham exposure.

### 2.3 Hind paw mechanical withdrawal threshold

Mice were acclimated to the testing room for two days prior to assessment of hind paw mechanical thresholds. On the day of testing, mice were placed in individual clear plastic chambers (11 x 5 x 3.5cm) on a 55cm-high wire mesh table and allowed to acclimate for 30 mins. The up-down method was performed using a series of monofilaments (1.65, 2.36, 2.83, 3.22, 3.61, 4.08, 4.31, 4.74 g; North Coast Medical, Inc Morgan Hill, CA, USA) applied to the right hind paw. The test started with application of the 3.22g monofilament. A negative response was followed by the next larger monofilament, whereas a positive response (brisk withdrawal of the paw) was followed by the next smaller monofilament. Four additional filaments were tested after the first positive response. The 50% withdraw threshold was calculated for each mouse and group means were determined as previously described [7].

### 2.4 Urinary bladder distention

Under inhaled isoflurane (4% induction, 2% maintenance), the bare ends of two Teflon-coated stainless-steel electrode wires (0.003″ diameter; Grass Technologies, West Warwick, RI) were implanted into the left and right abdominal muscle using a 26-gauge needle and the free ends were attached to a differential amplifier (Model 1700, A-M Systems, Sequim, WA). A 24-gauge angiocatheter (EXELINT, Los Angeles, CA, USA) was inserted into the bladder via the urethra and secured to the tail with tape. Isoflurane was reduced to approximately 1% until hind limb reflexes, but not escape behaviors, were present. Body temperature was maintained at approximately 37°C using a heating pad. The bladder was distended with compressed nitrogen gas controlled by a custom-made distension control device (The University of Iowa Medical Instruments, Iowa City, IA), manually adjusted by a dual-stage low delivery pressure regulator (Matheson-Linweld, Kansas City, MO), and verified by a separate pressure monitor (World Precision Instruments, Sarasota, FL). After stable responses to 60mmHg were confirmed, each pressure (15, 30, 45, 60mmHg) was applied in triplicate for 20 seconds with a 2-minute rest period in-between. Electromyographic activity was amplified, filtered, and recorded (Spike 2, Cambridge Electronic Design, Cambridge, UK) and the visceromotor response (VMR) was quantified and expressed as a percentage of baseline activity immediately prior to the distention.

### 2.5 Sucrose preference testing

Mice were housed individually with free access to two identical polysulfone drinking bottles (BioDAQ Liquid Choice Unplugged Allentown cages, Biological Data Acquisition, New Brunswick, NJ). Mice were acclimated to caging conditions for 24 hours with both bottles containing standard drinking water. One bottle was then filled with 1% or 2% sucrose and bottle weights were measured and bottle positions were interchanged daily. Volume and percentage of 1% or 2% sucrose was calculated for each mouse.

### 2.6 Nest building test

Mice were individually transferred to a clean cage containing no environmental enrichment outside of a 3.0 g nestlet square one hour before the start of the dark phase (5pm). Seventeen hours later, the nest and intact nestlet pieces were photographed and weighed. Two blinded experimenters scored the nests on a 1-5 scale according to previous publications [11; 12].

### 2.7 Serum corticosterone

Mice were deeply anesthetized with inhaled isoflurane (>5%) and trunk blood was collected. Blood was allowed to clot on ice for 1 hour and then centrifuged at 10,000 rpm for 10 minutes. Serum (clear supernatant) was collected and stored at −20°C until analysis. Serum corticosterone (CORT) was quantified using an ELISA kit according to the manufacturer’s instructions (ALPCO, Salem, NH).

### 2.8 mRNA extraction and qRT-PCR

Mice were deeply anesthetized with inhaled isoflurane (>5%) and bladder and whole brains were removed and snap frozen in liquid nitrogen or on dry ice, respectively. Hypothalamus and hippocampus were dissected, immediately snap frozen in liquid nitrogen, and stored at −80°C. All tissue was homogenized in Trizol reagent (Ambion, Austin, TX, USA), followed by manufacturer’s instructions of RNeasy Micro Kit (QIAGEN, Valencia, CA, USA). The concentration of RNA was determined by a NanoDrop™ 2000c Spectrophotometer (Thermo Fisher Scientific, Wilmington, DE, USA) and cDNA was made by using the iScript cDNA synthesis kit (Bio-Rad, Hercules, CA, USA). Finally, 5 μL of cDNA was mixed with SsoAdvanced SYBR Green Supermix (Bio-Rad) and indicated primers (Integrated DNA Technologies, Coralville, IA), and quantitative RT-PCR was performed using a Bio-Rad CFX manager 3.1 real time PCR system. GAPDH and b-actin were used as control genes for brain and bladder tissue, respectively. Samples were run in triplicate with negative controls. Efficiency values were derived for each individual sample (LinRegPOCR software v2012.3) and threshold cycle values were subtracted from those of the control gene and percentage of fold change over sham was calculated using the Pfaffl method [41].

### 2.9 Statistical analysis

Statistical analyses were performed using Two-way analysis of variance (ANOVA), with or without repeated measures, followed by Bonferroni’s posttest or Fisher’s least squared difference, as denoted in the manuscript (IBM SPSS Statistics v. 24, IBM Corporation, Armonk, NY; GraphPad Prism 8, GraphPad Software, Inc., La Jolla, CA). All data are expressed as mean ± SEM and *p* < 0.05 was considered significant.

## 3. Results

### 3.1 Foot shock-induced acute and persistent bladder hypersensitivity and hind paw allodynia in female A/J mice

Female A/J mice were exposed to 10 days of foot shock treatment and then tested for bladder sensitivity 1 or 28 days later. Compared to sham-exposed mice, shock-treated mice had a significant increase in the visceromotor response (VMR) during urinary bladder distension (UBD) 1 day after the last shock treatment (Figure 1A). This was also observed for 30mmHg and 60mmHg pressures at 28 days after the final shock treatment (Figure 1B). Comparisons of the area under the curve for the VMR at both time points revealed a significant effect of foot shock exposure, especially at 28 days post-shock (Figure 1C).

**Figure 1.**
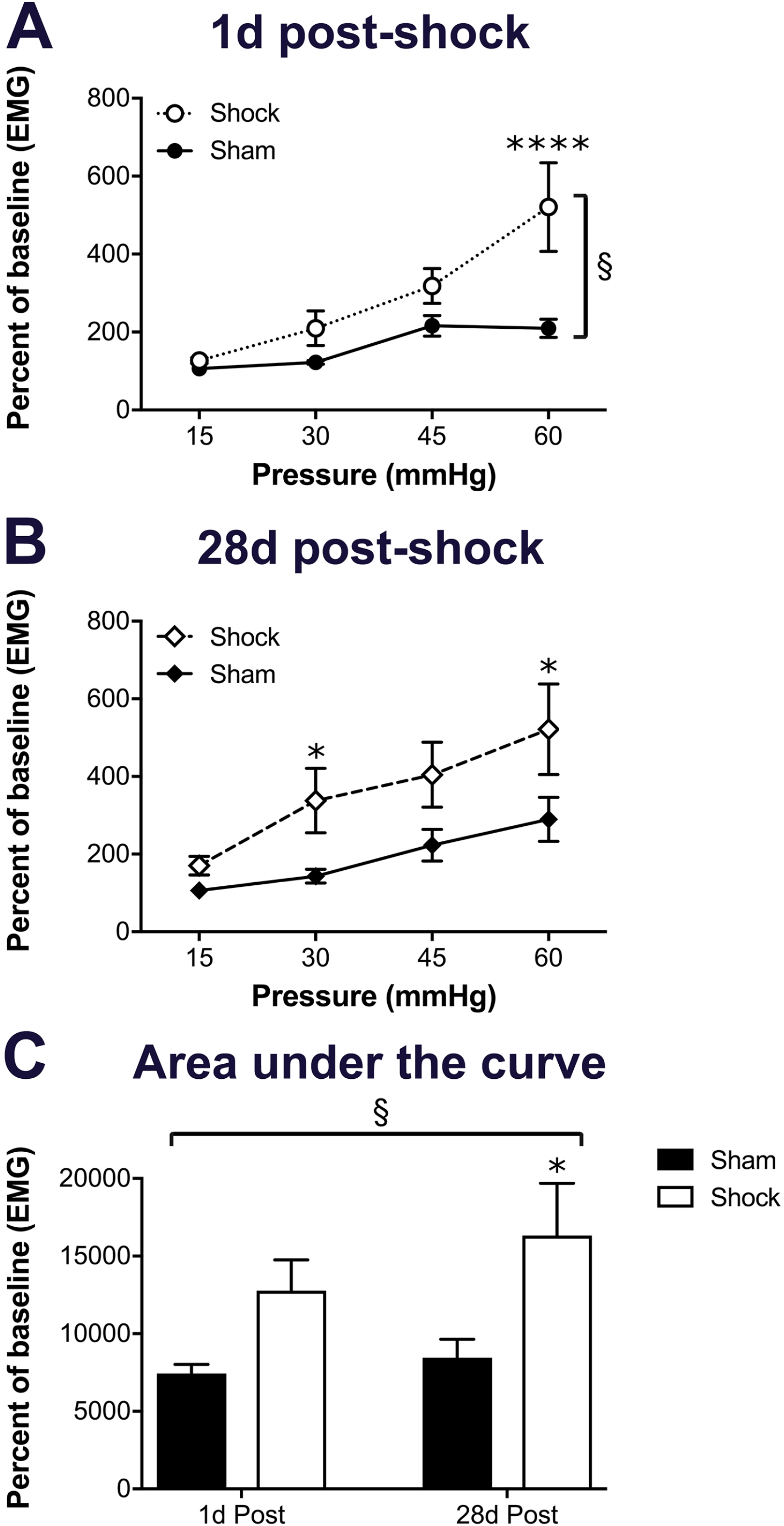
The visceromotor response (VMR) during urinary bladder distension (UBD) was measured 1d and 28d after the final foot shock exposure. A) At 1d post-foot shock, there was a significant impact of foot shock, especially at the 60mmHg pressure. B) At 28d post-foot shock, the shock group had a significantly higher VMR at 30mmHg and 60mmHg. C) Area under the curve (AUC) measurements revealed a significant impact of foot shock, particularly in the 28d group compared to sham. Brackets indicate a significant effect of shock (§ *p*<0.05), two-way RM ANOVA (A-B) or two-way ANOVA (C); *, **** *p*<0.05, 0.0001 vs. sham, Bonferroni posttest. n=5-8 for all groups.

Hind paw mechanical withdrawal thresholds were also measured at 1- and 28-days post-foot shock exposure. Shock-exposed mice had significantly lower withdrawal thresholds than their baseline measurements or those of sham-exposed mice, at 1-day post-foot shock exposure (Figure 2A). At 28 days post-foot shock, the foot shock-exposed mice maintained a significant reduction in withdrawal threshold, compared to their own baseline measurements, but were not significantly different from sham-exposed mice (Figure 2B).

**Figure 2.**
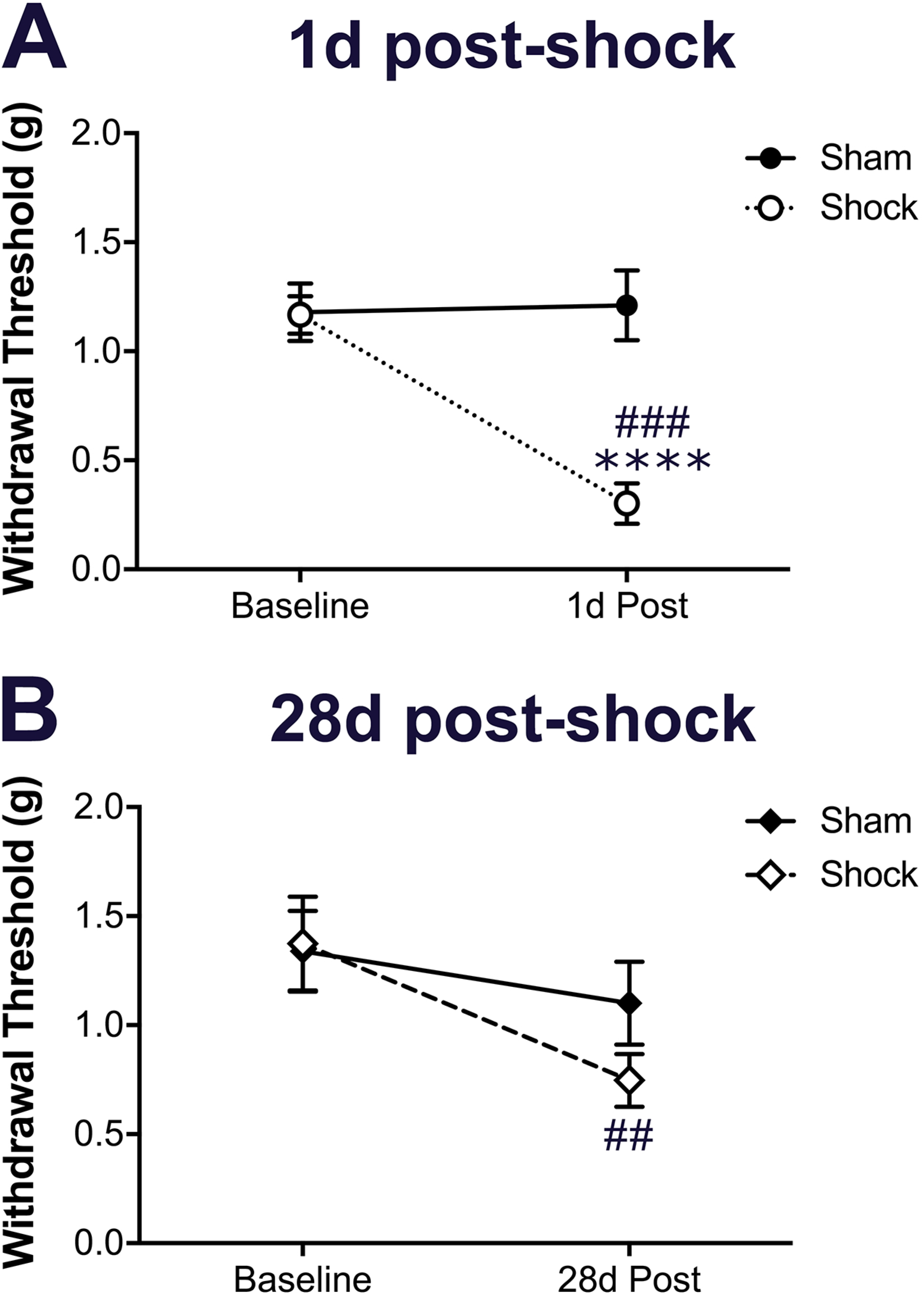
Hind paw mechanical withdrawal thresholds were measured 1d and 28d after the final foot shock exposure. A) The foot shock-exposed group had significantly lower mechanical withdrawal thresholds at 1d compared to both their own baseline measurements and the sham-exposed group. B) The foot shock-exposed group maintained significantly decreased mechanical withdrawal thresholds at 28d post-foot shock, compared to their baseline measurements, but were not significantly different from sham-exposed thresholds. Two-way RM ANOVA; **** *p*<0.05, 0.0001 vs. sham, ##, ### *p*<0.01, 0.001 vs. baseline, Bonferroni posttest. n=8 for all groups.

### 3.2 Depression-like behavior during and after foot shock exposure

Anhedonia was assessed both during and after foot shock exposure to determine depression-like outcomes related to stress in female A/J mice. Preference for 1% or 2% sucrose was measured for four days prior to and continually during the 10 days of foot shock exposure. Compared to sham-exposed mice, the percentage of 1% sucrose consumed by foot shock-exposed mice decreased significantly during 10 days foot shock, particularly on days 3-4 and 9-10 (Figure 3A). In contrast, preference for 2% sucrose was not significantly impacted by foot shock stress; however, there was a significant variance due to time across both groups (Figure 3B).

Nest building was assessed to evaluate anhedonia at 1- and 28-days post-foot shock exposure. Nest scores and the weight of remaining intact nestlet were both significantly impacted by foot shock and time (Figure 3C-D). Although the 1- and 28-day measurements were significantly different from their sham-exposed counterparts, the 28-day measurements had slightly improved and were significantly different from the 1-day measures (Figure 3C-D).

**Figure 3.**
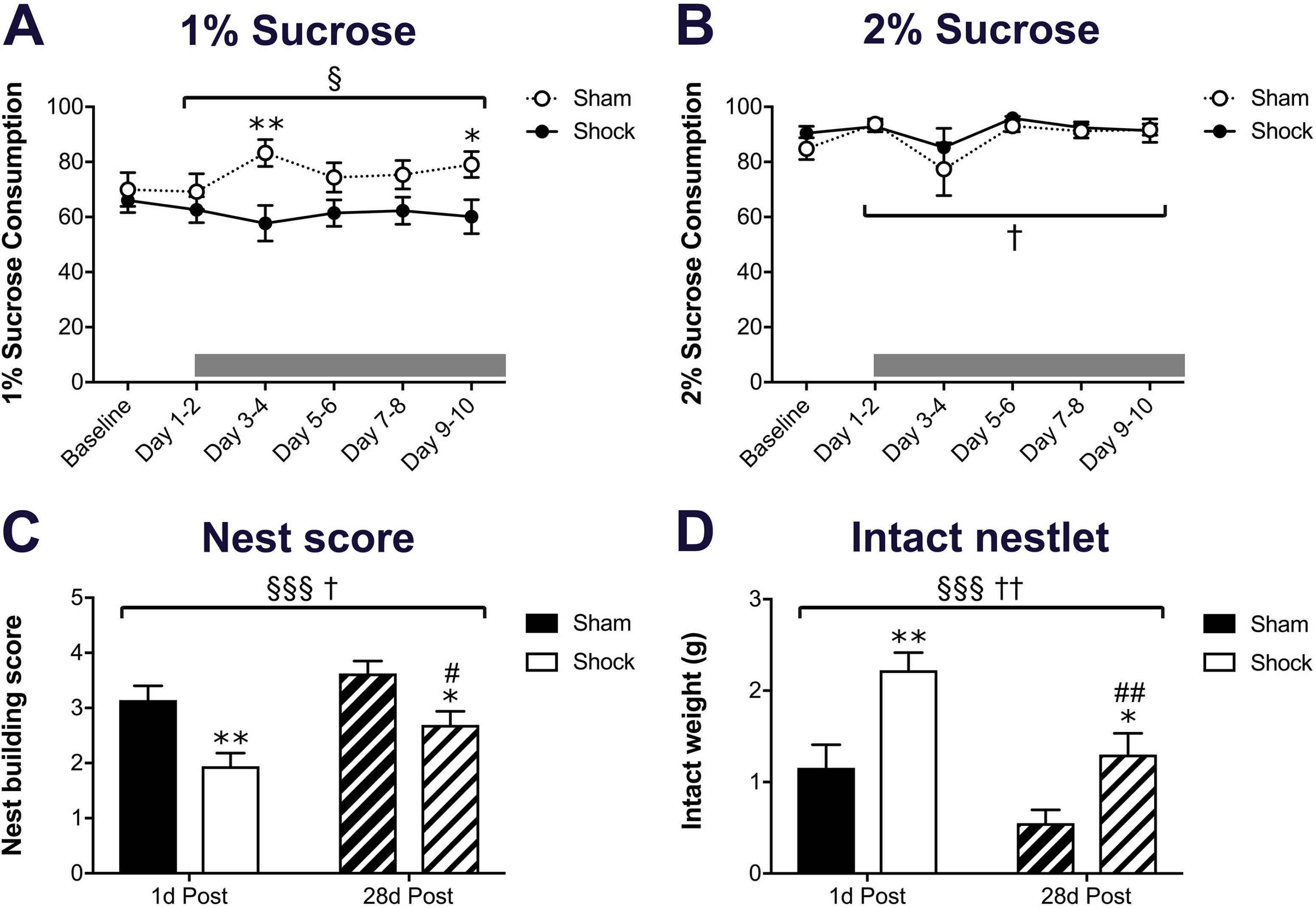
Sucrose preference testing and nest building were performed to assess anhedonic behavior 1d and 28d after the final foot shock exposure. Preference for 1% (A) and 2% (B) sucrose was measured prior to and during the 10 days of foot shock exposure. A) The percentage of 1% sucrose that was consumed, compared to standard drinking water, was significantly impacted by foot shock, particularly on days 3-4 and 9-10. B) The percentage of 2% sucrose that was consumed was significantly impacted over time, but not by foot shock exposure. Nest quality (C) and the weight of intact nesting material (D) were both significantly impacted by foot shock exposure and time. At 1d and 28d, both measurements were significantly different between sham- and foot shock-exposed mice. The 28d measurements in foot shock-exposed mice were significantly different from 1d. Brackets indicate a significant effect of shock (§, §§§ *p*<0.05, 0.001) or time (†, †† *p*<0.05, 0.01), two-way RM ANOVA (A-B) or two-way ANOVA (C-D); *, ** *p*<0.05, 0.01 vs. sham, #, ## *p*<0.05, 0.01 vs. baseline, Bonferroni posttest. n=7-8 for all groups.

### 3.3 Foot shock alters expression of selected mRNAs in the bladder but not mast cell infiltration or degranulation

Previous studies revealed that early life stress exposure increased mast cell degranulation and pro-inflammatory cytokine and growth factor expression in the bladder, which may contribute to urogenital hypersensitivity [42]. Therefore, we investigated the effect of foot shock exposure on mast cell infiltration and degranulation and levels of selected mRNAs in the bladder. There was no difference in the number of infiltrated mast cells or in the mast cell degranulation rate between bladders from 1-day post-foot shock and sham exposed mice (Figure 4A). A significant impact of foot shock or time was observed for every mRNA evaluated. The mRNA levels of interleukin-10 (IL-10), an anti-inflammatory cytokine, were significantly decreased by foot shock exposure, particularly at the 1-day time point (Figure 4D). The mRNA levels of both artemin and stem cell factor (SCF) were significantly elevated 28 days after foot shock exposure compared to 1 day, driving an overall time effect for both mRNAs (Figure 2D). Finally, levels of mRNA encoding monocyte chemoattractant protein 1 (MCP-1) were significantly decreased by foot shock exposure, particularly at the 28-day time point (Figure 4D).

**Figure 4.**
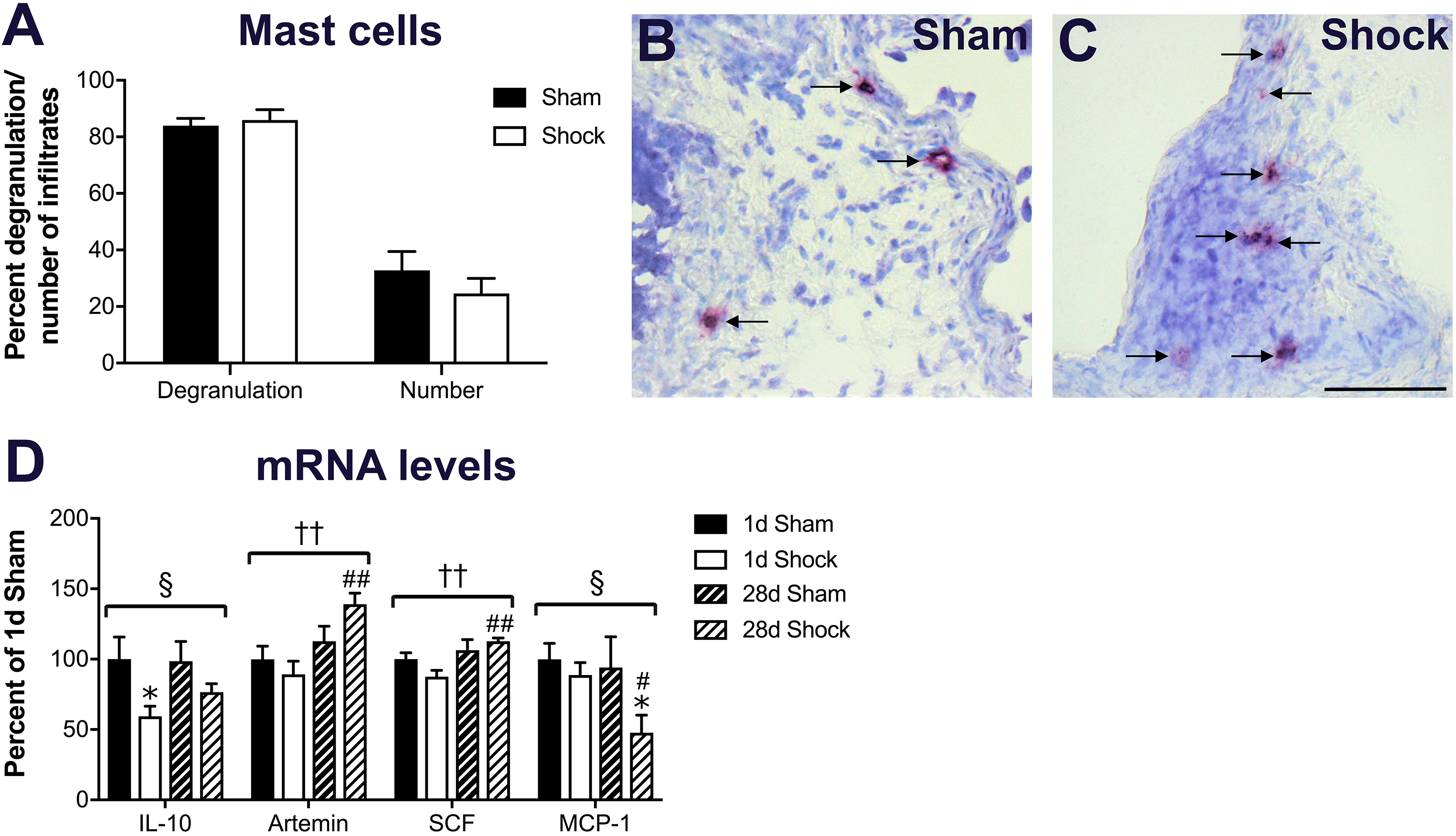
Bladder mast cell infiltration and degranulation, along with inflammatory-related mRNAs, were measured after foot shock exposure. Mast cells were visualized in the bladder using acidified toluidine blue in both sham- (B) and foot shock-exposed (C) mice. Nearly all mast cells displayed some degree of degranulation, evidenced as faint metachromasia and free granules within the cytoplasm and extruded outside of the cell borders (arrows). A) At 1d, there were no differences in the percentage of degranulation or the number of mast cells in the bladders between sham- or foot shock-exposed mice. D) The level of IL-10 mRNA was significantly impacted by foot shock exposure, particularly at the 1d time point. Artemin and SCF mRNA levels were similarly impacted over time with 28d foot shock bladders having significantly higher levels than 1d bladders. MCP-1 mRNA levels were also significantly impacted by foot shock, with 28d foot shock bladder having significantly lower levels than 28d sham or 1d foot shock bladder. Brackets indicate a significant effect of shock (§ *p*<0.05) or time (†† *p*<0.01), two-way ANOVA; * *p*<0.05 vs. sham, #, ## *p*<0.05, 0.01 vs. 1d Shock, Bonferroni posttest. n=6-8 for all groups.

### 3.4 Foot shock exposure transiently increases HPA axis output and regulation

To understand how foot shock stress impacts the output and regulation of the HPA axis in female A/J mice, we measured serum CORT and levels of selected mRNAs in hypothalamus and hippocampus. Serum CORT was significantly increased 1 day after completion of the foot shock paradigm compared to sham exposed mice (Figure 5A). Serum CORT levels in sham- and shock-exposed mice were not different 28 days later (Figure 5A). A significant overall effect of time was observed for mRNAs encoding both brain-derived neurotrophic factor (BDNF) and mineralocorticoid receptor (MR) in the hypothalamus, largely driven by a trend toward increased levels at 1-day post-shock that were significantly lower in 28-day post-shock samples (Figure 5B). In the hippocampus, glucocorticoid receptor (GR) mRNA levels were significantly higher in the sham group at 28 days compared to 1-day post-sham and 28 days post-shock (Figure 5C).

**Figure 5.**
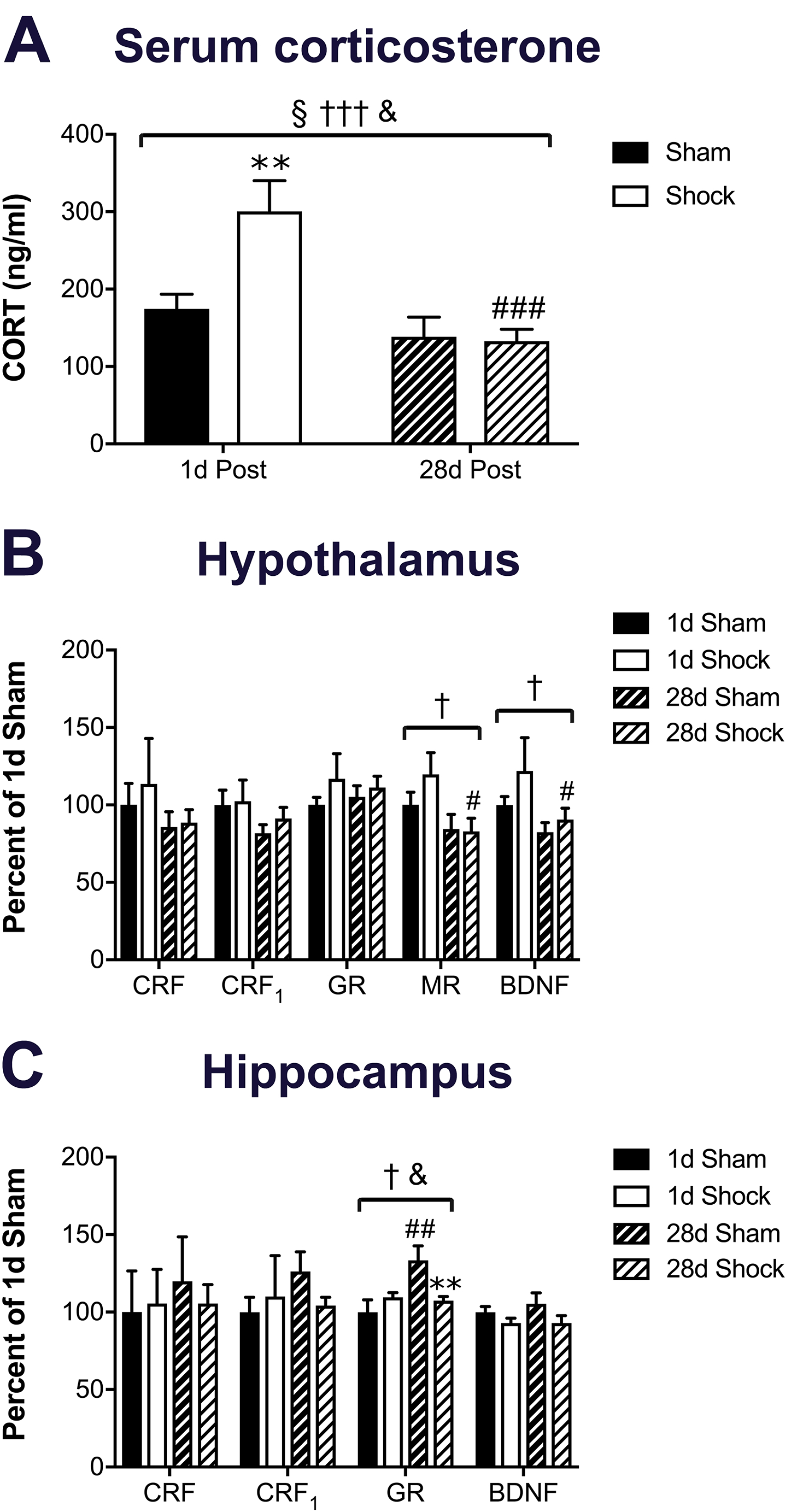
The output and regulation of the HPA axis was measured 1d and 28d after foot shock exposure. A) Serum corticosterone was significantly impacted by shock and/or time, such that 1d post-foot shock levels were significantly higher than 1d sham and 28d shock groups. B) Only mineralocorticoid receptor (MR) and brain-derived neurotrophic factor (BDNF) mRNA levels were impacted by foot shock in the hypothalamus, with the 28d shock groups having significantly lower levels of both mRNAs compared to 1d shock. C) A significant impact of time and a foot shock/time interaction effect was observed only for glucocorticoid receptor (GR) mRNA levels in the hippocampus. GR mRNA levels were significantly higher in 28d sham compared to 1d sham or 28d shock bladder. Brackets indicate a significant effect of shock (§ *p*<0.05), time (†, ††† *p*<0.05, 0.001), or a shock/time interaction (&, *p*<0.05), two-way ANOVA; ** *p*<0.01 vs. sham, #, ##, ### *p*<0.05, 0.01, 0.001 vs. 1d, Bonferroni posttest. n=4-8 for all groups.

## Discussion (1500 words)

It is well-established that stress exacerbates symptoms of chronic pain and that anxiety is a common comorbidity with widespread pain [8; 29; 32]. The influence of stressors on bladder sensitivity has been explored in rat models with genetically enhanced [30; 47] or comparatively normal [48] levels of behavioral anxiety. Here we have investigated A/J mice, an inbred mouse strain that exhibits comparatively high levels of anxiety, for bladder-specific and more widespread hypersensitivity, as well as for measures of depression and regulation of the HPA axis following a 10 day-long exposure to foot shock stress.

Strain differences in behaviors related to anxiety, depression, and pain have been widely reported for both mice and rats. A comparison of C57BL/6J and A/J mice in a series of anxiety-provoking behavioral tests showed that A/J mice of both sexes scored higher than C57BL/6J in all tests performed [28]. A/J mice were also one of the more anxiety-prone strains in many applied behavioral tests, such as open field, light/dark transition test, and elevated plus maze when compared to 13 other inbred strains (e.g. BALB/cJ, NIH/Nude, C3H/HeN, DBA/2J, AKR/J, C57BL/6ByJ, C57BL/10J) [9]. Significant strain effects on visceral sensitivity and behavioral measures of anxiety and depression were reported and most pronounced in the CBA/J strain from Harlan, which is derived from the same progenitor line as the A/J mice used in this study, albeit with multiple decades of genetic drift. [36]. Finally, Wistar-Kyoto rats have been identified as an anxiety-prone strain and display significant defeat behaviors during forced swim [2] and sex-specific anxiety-like behaviors, depending on the applied test [6].

Our paradigm of repeated exposure to foot shock stress in an anxiety-prone strain of mice in the current study paralleled previous studies incorporating stress to exacerbate hypersensitivity in anxiety-prone rat strains [30; 47; 49]. Robbins, et al. [47] demonstrated that chronic (10 days), but not acute (1 day), exposure to water avoidance stress increased VMR during UBD in Wistar-Kyoto but not Sprague Dawley rats. Additional studies have demonstrated that Wistar-Kyoto rats develop both acute and persistent bladder hyperalgesia, lasting up to 61 days [30], and altered micturition patterns [49] following 10 days of water avoidance stress exposure. Here, we observed a significant increase in bladder sensitivity both at 1 day and 28 days after 10 days of foot shock exposure. Chronic (7 days) exposure to foot shock has been shown to acutely increase bladder sensitivity in Sprague Dawley rats [48]; however, the duration of the increase was not reported, making it difficult to determine the influence of the strain on chronicity of hypersensitivity. Regardless, it is likely that genetic signatures in anxiety-prone strains make them more susceptible to outcomes related to chronic stress exposure, including long-lasting visceral hypersensitivity.

The A/J mice also displayed acute and persistent hind paw mechanical allodynia following 10 days of foot shock exposure. A sustained increase in hind paw mechanical withdrawal responses to a non-noxious stimulus was also reported by Lee et al., [30] following chronic water avoidance stress in Wistar-Kyoto rats. Interestingly, a single exposure to water avoidance stress generated both mechanical and thermal hind paw analgesia in Wistar-Kyoto rats, while a 10-day exposure had no effect compared to sham-exposed controls [47]. Clinically, IC/PBS patients with widespread pain report sleep disturbances, increased depression and anxiety scores, increased somatic symptom burden, and greater negative affect scores, compared to IC/PBS patients with pain localized only to the pelvis or abdominal region [29]. It is hypothesized that patients with widespread sensitivity and high comorbidity have a centralized pain phenotype that is more susceptible to acute and chronic stress exposure [15; 29]. This is largely based on functional magnetic resonance imaging (fMRI) and magnetic resonance spectroscopy (MRS) studies of patients within the same study cohort showing increased gray matter volumes and functional connectivity between the somatosensory and insular cortices [27], which was also related to higher choline levels in the anterior cingulate cortex (ACC) [19], all of which correlated with negative mood. Interestingly, Wistar-Kyoto rats exposed to chronic water avoidance stress exhibited greater cortical activation in many of these same areas in response to passive bladder distention [55], supporting the use of chronic stress in anxiety-prone rodents to model UCPPS.

Sucrose preference is a commonly used method with high validity for evaluating reward behavior, as preference is decreased or increased following stress exposure or antidepressant treatment, respectively [45]. Indeed, in this study, we observed a significant decrease in preference for a 1% sucrose solution during the duration of the foot shock exposure. Previous studies have investigated genetic contributions to sucrose preference and have identified variants in relevant taste genes, particularly Tas1r3 [3]. A specific deleterious mutation in Tas1r3 was identified in the low sucrose-preferring 129X1/J strain, as well as in four other strains including A/J, that was not present in the high sucrose-preferring C57Bl/6 strain [46]. However, in a comparison study of 11 different strains using 9 concentrations of sucrose, the A/J strain was one of the highest sucrose-consuming strains tested, suggesting that sucrose preference is under polygenic control [31]. Our reported ~70% preference for a 1% sucrose solution is similar to that reported in A/J mice by Lewis et al. [31], and the foot shock-induced decrease in preference suggests a reduction in reward-seeking behavior. The study by Lewis et al., [31] also reported a lack of correlation between preference of different concentrations, as well as a compensatory effect of reduced caloric intake from chow as sucrose intake increased, particularly in the A/J strain. This suggests that there may be increasing metabolic influences regulating sucrose intake at higher concentrations, making these observations less dependent upon reward-based behaviors, which may underlie the lack of foot shock-induced reduced preference for 2% sucrose in the current study.

Nest building is a complex and innate behavior that is negatively impacted by hippocampal damage [13], and has been used to assess rodent models of pain disorders, schizophrenia, and neurodegenerative disease [24]. Nest building scores were lower in a model of social defeat stress and reversed by select anti-depressant treatments [39], suggesting that it is an appropriate measure for depressive-like behaviors. Here, we observed a persistent decrease in nest quality following exposure to foot shock. The decreased nest building is likely more indicative of general distress rather than persistent pain, as a previous study showed that carprofen analgesia had no impact on nest building in a model of post-surgical pain [25]. The persistence of anhedonic behavior, coinciding with the increased bladder and hind paw sensitivity, supports the validity of this model for the study of stress-induced, widespread hypersensitivity.

The role of CRF in influencing pain signaling has been well-described in both the peripheral and central nervous system. Peripherally-released CRF activates mast cells [52] and binds receptors expressed by urothelium [18], which in turn induce neurogenic inflammation and urothelial leakiness, respectively. Here, we showed no increase in mast cell infiltration or degranulation following foot shock exposure; however, it should be noted that the level of degranulation for both sham- and shock-exposed mast cells in the bladder was markedly high (>80%). This is much higher than we reported in C57Bl/6 bladder (~30%) [42] and is similar to reports of the high anxiety strain of Wistar-Kyoto (WKY) rats (~75%), which also showed no increase in degranulation after a 10-day water avoidance stress protocol [30; 40; 50]. The comparatively high level of serum CORT in the sham cohorts (~170ng/ml in A/J vs. 20ng/ml in C57Bl/6 [43]) suggests that over-activity within, or downstream from, the HPA axis may be driving increased mast cell degranulation in A/J mice. Increased NGF and pro-inflammatory cytokine levels were observed in sera from IC/PBS patients [23] and from cold cup biopsies, which also had an increase in mast cell infiltration [33]. Here, we observed an overall pro-inflammatory shift in gene expression in the bladder with a decrease in IL-10 and an increase in both artemin and SCF mRNA levels. The persistent increase in artemin mRNA levels mimics other rodent studies of inflammation- or nerve injury-induced persistent pain [35] [21]. Pre-clinical studies have implicated SCF and the kit pathway in overactive bladder [26; 38]. MCP-1 has been reported to play an important role in inflammatory-related diseases such as atherosclerosis and rheumatoid arthritis [5; 51], and MCP-1 levels are higher in urine and bladder from rodent models of IC/PBS [4; 34].Our observation of significantly decreased MCP-1 mRNA levels at 28 days after foot shock was somewhat surprising and may represent a compensatory response.

In conclusion, exposure to foot shock stress elicited prolonged widespread hypersensitivity in the anxiety-prone A/J mouse strain, evidenced as increased bladder sensitivity and decreased hind paw mechanical withdrawal thresholds. Anhedonia was also observed as evidenced by decreased sucrose preference during the foot shock exposure and poor nest quality. A prolonged shift towards a pro-inflammatory gene expression profile in the bladder occurred following a transient increase in serum CORT levels. Together, these data suggest that A/J mice may provide an excellent model to understand the mechanisms contributing to widespreadness of pain and increased comorbidity in a subset of UCPPS patients.

**Table 1.**
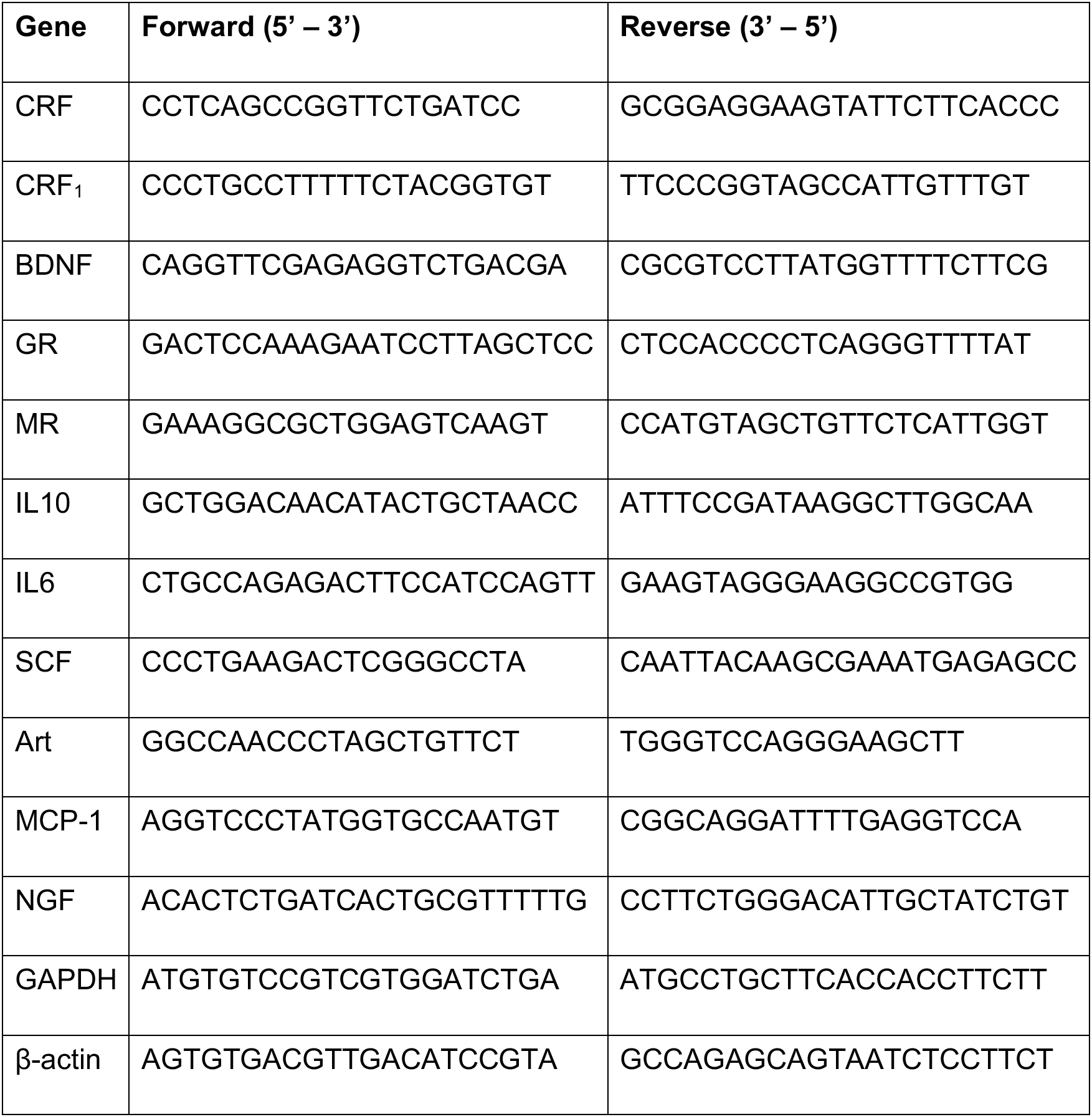
Primers used for real-time PCR analysis

## Acknowledgments

The authors would like to thank Dr. Isabella Fuentes, Ruipeng Wang, and Janelle Ryals for technical assistance and Drs. Ken McCarson, Andrea Nicol, and John Thyfault for expert advice and feedback on experimental design, methodology, and interpretation.

## Competing Interests

None of the authors of this work have competing or conflicting interests in the associated work.

## Funding

This work was supported by National Institutes of Health (NIH) grants R01DK099611 (JAC), R01DK103872 (JAC), R01NS043314 (DEW), P20 GM103418 from the Idea Network of Biomedical Research Excellence (INBRE) Program (DEW,), and the Kansas IDDRC, U54 HD 090216.

## Author Contributions

PYW directed, designed, and performed the experiments and was the principle author of the manuscript. PYW, XW, JAC, and DEW were involved in specific aspects of data collection, analysis, and manuscript edits. JAC and DEW helped design all experiments with PYW and were principle authors of the manuscript.

